# Ecophysiological behaviour of major *Fusarium* species in response to combinations of temperature and water activity constraints

**DOI:** 10.1101/2024.09.19.613940

**Authors:** Marie-Anne Garcia, Rémi Mahmoud, Marie-Odile Bancal, Pierre Bancal, Stéphane Bernillon, Laetitia Pinson-Gadais, Florence Richard-Forget, Marie Foulongne-Oriol

## Abstract

*Fusarium* Head Blight (FHB) is a devastating fungal disease affecting cereals, caused by *Fusarium* species that can produce harmful mycotoxins. *Fusarium* species share the same ecological niche, and their population dynamic and associated mycotoxin patterns are driven by the environment. The aim of the present study was to investigate ecophysiological characteristics of the major *Fusarium* species causing FHB under abiotic factors. Growth and mycotoxin production of different strains of *Fusarium avenaceum, Fusarium graminearum*, *Fusarium langsethiae, Fusarium poae* and *Fusarium tricinctum* were characterized under a combined effect of temperature (ϴ = 15, 20, 25 and 30°C) and water activity (a_w_ = 0.99, 0.98, 0.97, 0.96, 0.95 and 0.94). Using innovative statistical analyses, we demonstrated that those *Fusarium* species greatly differ in their responses to the studied environmental constraints. Our findings indicated that ϴ, a_w_, and their interaction were the major factors with a significant impact on the species behaviour. The intraspecific variation was demonstrated as less pronounced than the interspecific one. Understanding the ecophysiological requirements of *Fusarium* species is crucial in the context of climate change that is predicted to worsen disease outbreaks. Our data constitute a valuable knowledge base for improving the reliability and robustness of FHB prediction models and anticipating the associated mycotoxin risk.

**IMPORTANCE:** *Fusarium* species pose a significant threat to major cereal crops, including wheat. These fungi not only reduce yield but also produce mycotoxins harmful to animals and humans. The prevalence of each *Fusarium* species is influenced by environmental conditions and shifts in pathogen populations, leading to changes in mycotoxin patterns have already been observed in relation to climate changes. This study revealed distinct ecophysiological behaviours, including mycelium growth and mycotoxin production, among the five major *Fusarium* species when exposed to varying temperature and water activity conditions. Our findings provide a valuable foundation for a more comprehensive understanding of the challenge of mycotoxin contamination and for the development of more effective mitigation strategies in the near future.

## INTRODUCTION

*Fusarium* Head Blight (FHB) is a devastating fungal disease affecting small grain cereals worldwide. FHB is mainly caused by several *Fusarium* species that can produce various mycotoxins harmful to humans and animals. In addition to economic losses due to reduced yield and a lower grain quality, mycotoxin-producing *Fusarium* species also induce important waste due to the downgrading of contaminated batches. Among the species causing FHB in Europe, *Fusarium graminearum* is acknowledged as the key actor. But, *F. graminearum* is never found alone and commonly co-occurs with other *Fusarium* species during infection. *Fusarium avenaceum* and *Fusarium tricinctum* are frequently found associated with FHB (Xu *et al*. 2005; Beccari *et al*. 2018; Vogelgsang *et al*. 2019; Valverde-Bogantes *et al*. 2020; Infantino *et al*. 2023; Dubois and Roucou 2023). Recent works also highlighted the increased occurrence of other species in Europe such as *Fusarium poae* or *Fusarium langsethiae* (Osborne and Stein 2007; Valverde-Bogantes *et al*. 2020; Infantino *et al*. 2023; Dubois and Roucou 2023). Each *Fusarium* species has its own mycotoxin profile. *F. graminearum* is a well-known producer of type B trichothecenes (TCTB) including DON and its acetylated derivates (3-acetyldeoxynivalenol (3-ADON) and 15-ADON), nivalenol (NIV) and of zearalenone (ZEA) (Van Der Lee *et al*. 2015). *F. langsethiae* and *F. poae* produce type A trichothecenes (TCTA) including diacetoxyscirpenol (DAS), T-2 and HT-2 (Kokkonen *et al*. 2012; Dinolfo and Stenglein 2014) but *F. poae* can also produce TCTB (NIV and fusarenon-X (Fx)) as well as enniatins (ENN). *F. avenaceum* and *F. tricinctum* are enniatins (enniatins A (ENNA), ENNA1, ENNB and ENNB1) and beauvericin (BEA) producers (Gautier *et al*. 2020). All these *Fusarium* mycotoxins are known for their cytotoxic effect (Langseth *et al*. 1999), especially TCTB and TCTA, which are reported to be immunosuppressive and inhibit the synthesis of proteins in eukaryotes (Cundliffe *et al*. 1974; Ueno and Matsumoto 1975; Cundliffe and Davies 1977; Eudes *et al*. 2000; Steinmetz *et al*. 2009; Knutsen *et al*. 2017). ZEA is oestrogenic, *i.e.* it affects fertility and reproduction (Fink-Gremmels and Malekinejad 2007). The toxic effects of ENN and BEA are related to their ionophoric properties and therefore their capacity to affect the integrity of biological membranes (Mallebrera *et al*. 2018). In Europe, most of these mycotoxins are covered by regulations fixed by the European legislation. Maximum levels and guidance values for mycotoxins in cereals are constantly evolving due to new knowledge in toxicology and occurrence data but also to shifts in FHB pathogen populations associated with changes in mycotoxins contaminating crops (Pinotti *et al*. 2016).

Climate change is likely to deeply impact global food safety. One of the major concerns is the increasing risk of human exposure to mycotoxins (Garcia-Cela *et al*. 2018; Kos *et al*. 2023). Several studies have already demonstrated that climate change has led to substantial fluctuations in the distribution of FHB pathogens and the associated-mycotoxins profiles (Osborne and Stein 2007; Valverde-Bogantes *et al*. 2020). Changes in the prevalence of one *Fusarium* species over the others, together with the subsequent major mycotoxin contaminating harvest, are becoming more common in Europe (Beyer *et al*. 2014, Covarelli *et al*. 2015; Van Der Lee *et al*. 2015; Banik *et al*. 2018; Nogueira *et al*. 2018; Valverde-Bogantes *et al*. 2020). Understanding the biological and climatic factors influencing Fusarium populations is crucial for effectively managing FHB and mycotoxin risks

Previously mentioned studies indicated that, despite the ability of *Fusarium* species to occupy the same ecological niche and share the same biotic and abiotic environment, their intrinsic ecophysiological requirements could exhibit slight differences. This could potentially result in differential adaptive capacities in response to changing niche conditions, thereby driving the installation of the fittest species. In addition, *Fusarium* species are able to produce a large arsenal of secondary metabolites, including mycotoxins, which may confer competitive advantage during niche colonisation (Fox and Howlett 2008; Audenaert *et al*. 2013).

Several studies have investigated the *in vitro* responses of the predominant *Fusarium* species under abiotic constraints, mainly temperature and water activity, with a particular focus on growth and mycotoxin production (Hope *et al*. 2005; Nazari *et al*. 2014; Verheecke-Vaessen *et al*. 2022). Overall, those studies suggested that each species exhibits a distinctive response signature and that within a given species, optimal conditions required for mycelium growth and mycotoxin production are most often different. For example, 25°C is the optimal temperature for growth and DON production by *F. graminearum* (Hope *et al*. 2005), while *F. langsethiae* produce HT-2 and T-2 optimally between 15 and 35°C (Nazari *et al*. 2014). Knowledge about the way that abiotic parameters modulate fungal development and mycotoxin production of minor *Fusarium* species is much more limited (Gautier *et al. 2*020).Besides, the currently available data have been obtained using quite different experimental conditions (culture media or abiotic factors), mostly using only one or two strains per species which doesn’t allow to assess the adaptation capacity of the different species and speculate which species will be more suited to a particular environment.

To investigate the aforementioned knowledge gaps, the present work aimed to study the combined effect of environmental factors (temperature ϴ and water activity a_w_) on the growth of the five most frequently encountered *Fusarium* species (*F. avenaceum*, *F. graminearum*, *F. langsethiae*, *F. poae* and *F. tricinctum*) in European harvests and on their respective mycotoxin production. Twenty-four combinations of ϴ (15, 20, 25 and 30°C) and a_w_ (0.94, 0.95, 0.96, 0.97, 0.98 and 0.99) were tested and five strains of each *Fusarium* species were considered. The probability of growth was assessed for each *Fusarium* species, and calibrated logistic growth curves tailored to each species and strain were calculated. The impact of different factors (ϴ, a_w_, inter/intra-specific variability) on the growth’s curves parameters were investigated. Additionally, the same abiotic factors (a_w_ and ϴ) were tested for their effect on mycotoxin production which was considered as a binary response (presence or absence). Finally, correlations between growth parameters and mycotoxin production were studied.

The present study will provide essential insights that will allow anticipating the future shifts in *Fusarium* species distribution and mycotoxin patterns as a result of environmental changes.

## RESULTS

### INFLUENCE OF ENVIRONMENTAL CONDITIONS ON GROWTH

#### FREQUENCY OF GROWTH AND GROWTH PARAMETERS IN RESPONSE TO A_W_ AND ϴ FACTORS

Growth frequency (f_g_) and growth parameters (the time at the inflection point τ, the carriage capacity K, the relative growth rate r and the maximal growth rate Vmax) estimated under different a_w_ and ϴ variations for the five strains of the five studied *Fusarium* species are shown in Figure 1. Overall, data evidenced that, when growth occurred, the K, τ, r and Vmax parameters followed different patterns depending on the species, strains and the environmental factors (a_w_, ϴ).

**Figure 1:**
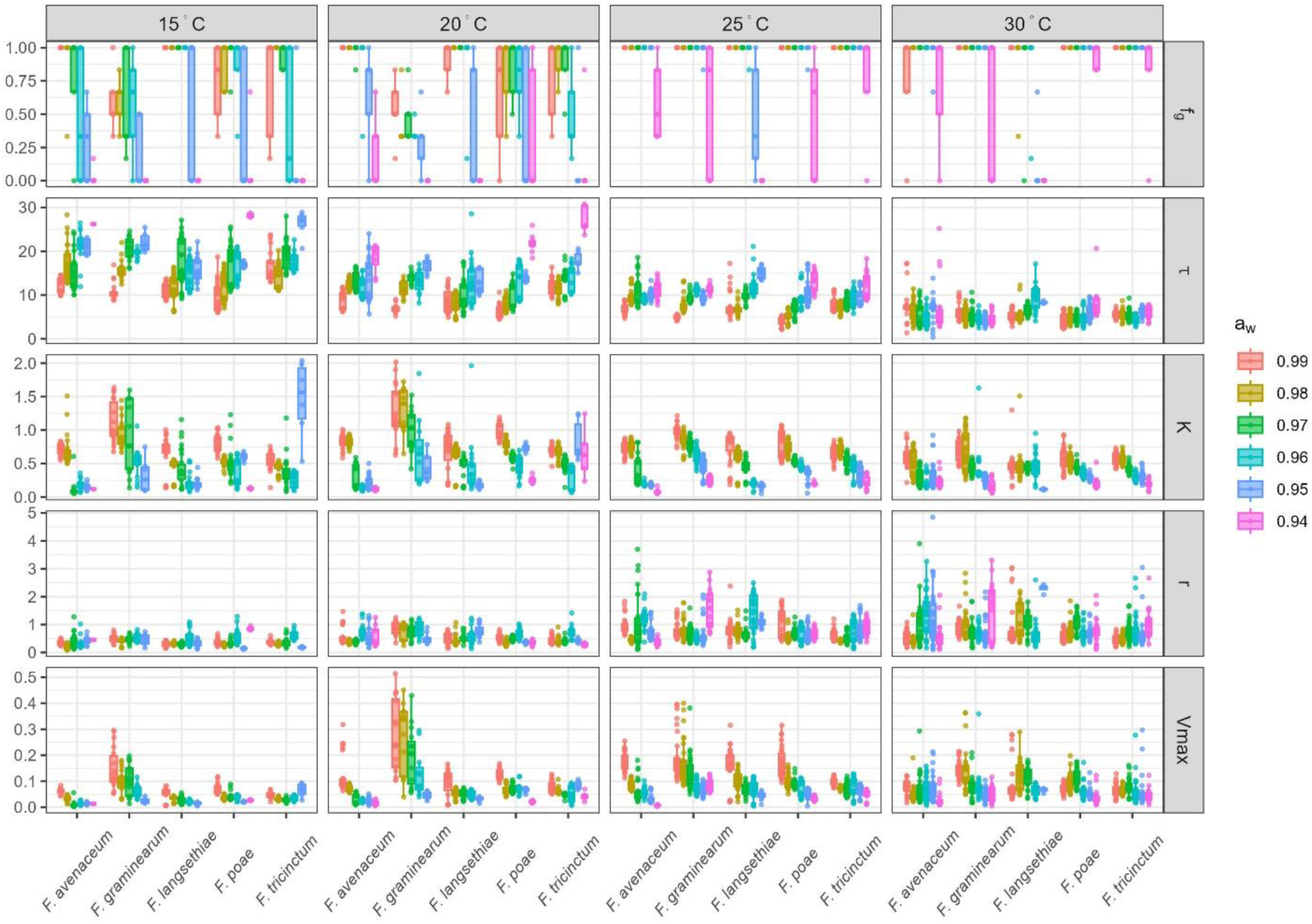
Growth frequency (f_g_) and growth parameters (τ, K, r and Vmax) for the five *Fusarium* species, according to environmental variation. Growth frequency (f_g_, expressed without unit) and growth parameters (τ expressed in days, K without unit, r and Vmax in OD_unit_.days^-1^) in lines, were estimated on 5 different strains per species (*F. avenaceum*, *F. graminearum*, *F. langsethiae*, *F. poae* and *F. tricinctum*, on the x-axis). Four different ϴ (15, 20, 25 and 30°C, showed in columns) and six levels of a_w_ (0.94, 0.95, 0.96, 0.97, 0.98 and 0.99) were tested. Each dot represents the mean frequency of the 6 replicates for each strain and the boxplots show the intraspecific diversity within the 5 species.

When considering first the growth frequency (f_g_), highest values (reached f_g_ = 1) were obtained at ϴ ≥ 25°C and a_w_ > 0.94 whatever the *Fusarium* species. Overall, fungal growth was occurring in 2672 out of the 3600 inoculated wells. Considering all species and conditions, *F. poae* and *F. avenaceum* exhibited the highest number of wells with detected fungal growth (80%), while *F. graminearum* was the least successful (66%). The highest heterogeneity in f_g_ was observed for the six a_w_ levels, at ϴ ≤ 20°C (Figure 1). Growth frequency (f_g_) data reported in Figure 1 also revealed the intraspecific variation of *Fusarium* in responses to environmental changes.

Concerning the τ_mean_ parameter, values determined for all studied *Fusarium* species and strains were, on average, the highest at 15°C (τ_mean_ = 15.8 days). These values gradually decreased when temperature increased to reach an average of 5.76 days at 30°C. On figure 1, it could also be noticed, that, with few exceptions, decreasing a_w_ led to an increased in τ_mean._. This pattern was particularly apparent for the 15°C, 20°C and 25°C conditions.

For most of the studied *Fusarium* species, K values were highest at 20°C or 25°C (Figure 1). The highest K values were observed for *F. graminearum.* Overall, K was shown to decrease with aw. For example at ϴ = 25°C, a four-fold decrease in the K value of *F. graminearum* was observed when a_w_ decreased from 0.99 to 0.94 (K = 0.98, 0.83, 0.78, 0.62, 0.45 and 0.24 at a_w_ = 0.99, 0.98, 0.97, 0.96, 0.95 and 0.94, respectively; Figure 1). Surprisingly, very high K values were fitted for *F. tricinctum* at quite challenging environmental conditions, *i.e.* K = 1.46 at ϴ = 15°C, a_w_ = 0.95 and K = 0.66 and 0.84 at ϴ = 20°C, a_w_ = 0.94 and 0.95, respectively. However, such an observation has to be considered as a mathematical artefact since under the former conditions, *F. tricinctum* showed a delayed growth (τ > 25 days) exceeding the duration of the experiment. Indeed, the fitted K could not be seen as indicative of observed growth when the τ value was too high (τ > 25 days), as maximal OD was never reached during the experimentation (Figure 1, Figure 3).

Regarding the r parameter, its value significantly increased (p-value < 0.05) with ϴ (r_mean_ = 0.37, 0.53, 0.77 and 0.82 OD_unit_.days^-1^ (ODU.D^-1^) at 15, 20, 25 and 30°C, respectively; Figure 1). Moreover, for growth conditions with a_w_ < 0.97, no significant difference between r parameter values determined for each *Fusarium* species was observed (p-value > 0.05).

Whatever the considered species, the Vmax parameter globally increased with ϴ, until 25°C, while a slight decrease was noted at 30°C (Vmax_mean_ = 90 and 75 mODU.D^-1^ at 25°C and 30°C, respectively). Among all the studied species, *F. graminearum* was the fastest-growing species with Vmax_mean_ = 97, 198, 122 and 92 mODU.D^-1^ at 15, 20, 25 and 30°C, respectively (Figure 1). On the contrary, *F. avenaceum* and *F. tricinctum* were the slowest-growing species with Vmax_mean_ = 74 and 72 mODU.D^-1^ at 25°C or Vmax_mean_ = 32 and 37 mODU.D^-1^ at 15°C, respectively. Overall, the Vmax parameter increased slightly with aw. Interestingly, at ϴ = 20°C and 25°C, the Vmax values assessed for *F. graminearum* showed a 4-fold and 2-fold decrease respectively between a_w_ 0.99 and 0.94.

#### PROBABILITY OF GROWTH : SPECIES EXHIBIT CONTRASTED RESPONSES TO A_W_ AND ϴ FACTORS

The five *Fusarium* species exhibited different growth responses to environmental variations. To investigate the specific effects of a_w_ and ϴ factors on growth parameters, mycelium growth was modelled. The probability model with the lowest Bayesian information criterion (BIC) reached an accuracy of 83%. As appeared in Figure 2, *F. langsethiae* displayed a distinct behaviour compared to the other species. Indeed, its growth probability (p_g_) strongly decreased at low a_w_ while minimally affected by ϴ. *F. langsethiae* appeared more adapted to lower temperature and higher water availability. On the other hand, p_g_ of the other species varied both with a_w_ and ϴ. For *F. graminearum, F. avenaceum, F. poae* and *F. tricinctum*, p_g_ exceeded 90% only when ϴ exceeded 25°C and a_w_ reached 0.97 (Figure 2). Moreover, *F. graminearum* appeared as the less robust; at 15°C and 20°C, its growth probability was lower than that of any other *Fusarium* species at all tested a_w_ levels.

**Figure 2:**
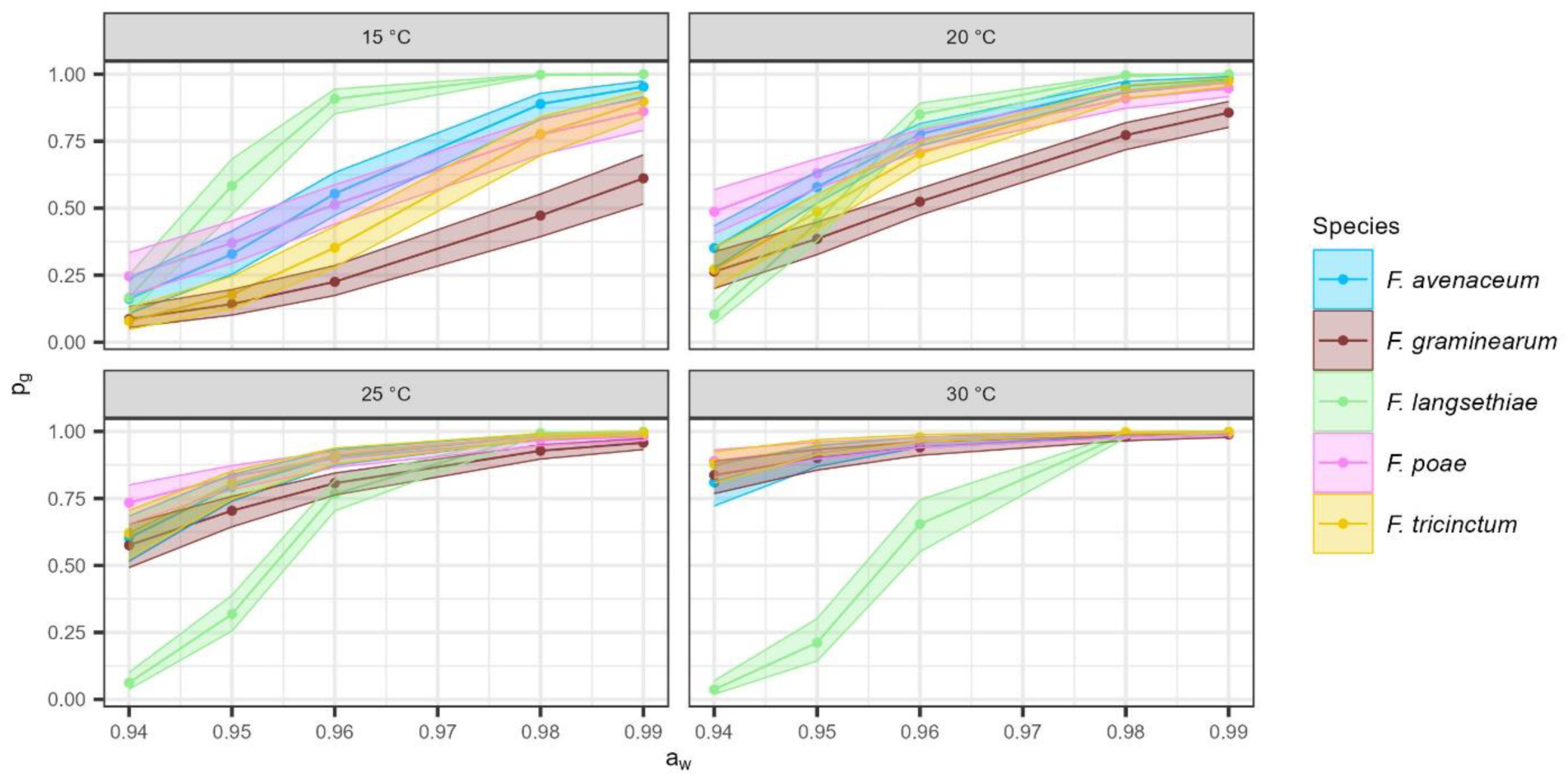
Modelling growth probability with logistic regression for the five *Fusarium* species, according to environmental variation. Growth probability (p_g_, on the y-axis) was estimated by the optimal logistic regression model (logit(p_g_) = β_0_ + (β_1_ + β_1i_)·ϴ_i_ + (β_2_ + β_2i_)·a_wi_ + β_3i_·fs_i_) against four ϴ (15, 20, 25 and 30°C, on each box) and at six levels of a_w_ (0.94, 0.95, 0.96, 0.97, 0.98 and 0.99; on the x-axis). Each dot represents the p_g_ of the five *Fusarium* strains for each species. Confidence intervals (95%) are represented by shading around the curves.

#### GROWTH PARAMETERS OF SPECIES ARE DIFFERENTIALLY DRIVEN BY A_W_ X ϴ CONDITIONS

Quadratic models, with the parameter as response variable and a_w_ and ϴ as covariates were applied for each studied *Fusarium* species and allowed us to establish response curves shown in Figure 3.

**Figure 3:**
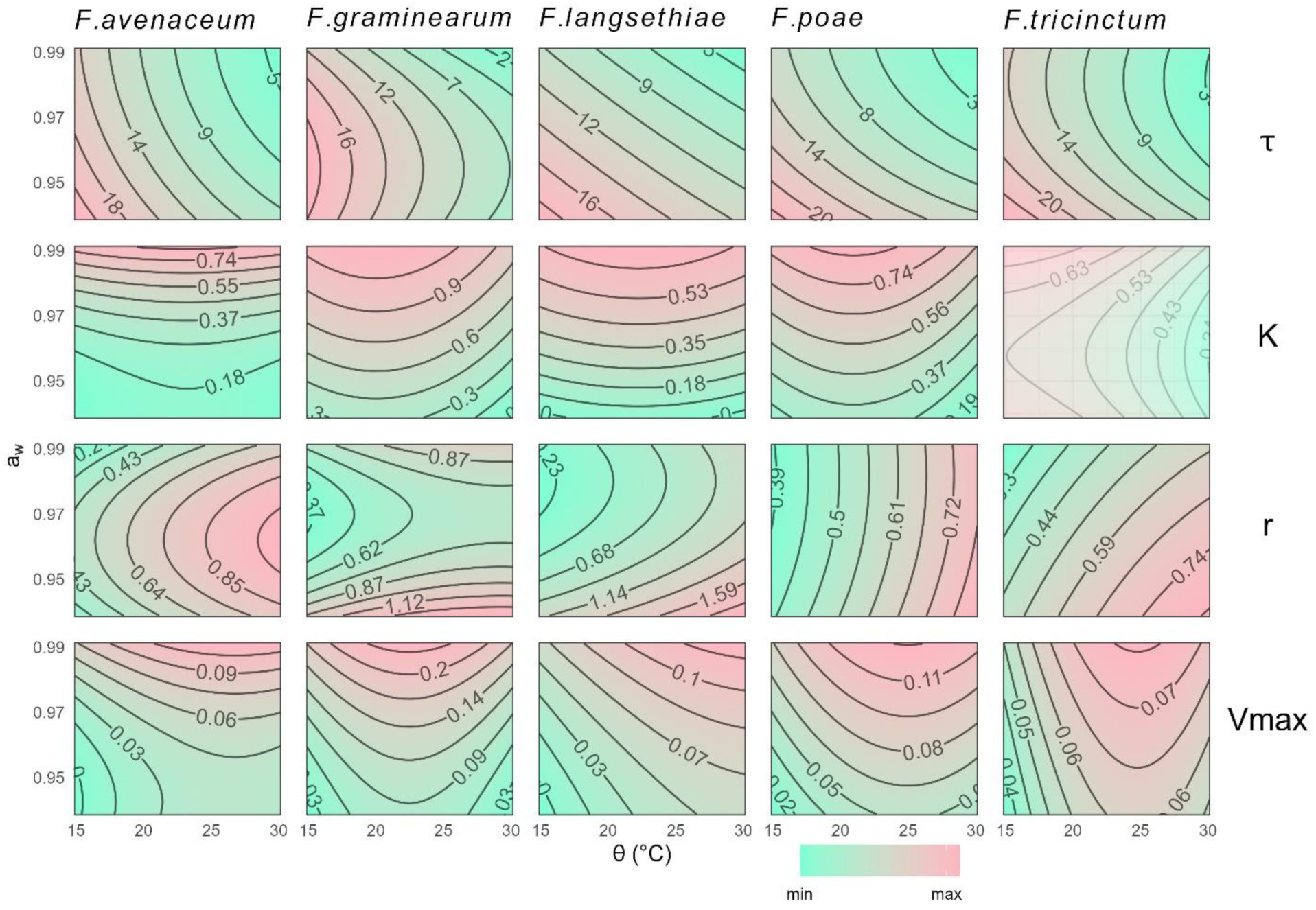
Response surface plots of growth parameters values for the five *Fusarium* species, according to combined abiotic conditions (a_w_ and ϴ). Contour plots of growth parameters (τ, K, r and Vmax in line) were estimated using quadratic models for each *Fusarium* species (*F. graminearum, F. avenaceum, F. langsethiae, F. poae* and *F. tricinctum* in column), as a function of a_w_ (y-axis) and ϴ (x-axis). Only grown replicates were considered. Each response surface has its own scale, this is indicated by numbers on the lines of the contour plot. The contour plot of K for *F. tricinctum* is shaded because the model did not fit correctly the response curves for this species.

Our data indicated a significant increase in time to reach the inflection point (τ) in response to lower temperature (ϴ ≤ 20°C) and reduced a_w_, regardless of the species (p-value < 0.05). Indeed, τ values were maximal at low ϴ, with little influence of either a_w_ or species (Figure 3).

In contrast, the carriage capacity (K) values were maximal at high a_w_ levels, with little or no influence of ϴ (Figure 3). The quadratic model was unable to correctly fit the K observed values for *F. tricinctum* because of the challenging estimation of this parameter for this species (as described above, in the “Growth parameters and frequency of *Fusarium* species in response to a_w_ and ϴ factors” section).

The relative (r) and maximum (Vmax) growth rates also exhibited a different response to a_w_ and ϴ depending on the fungal species (Figure 3). The r parameter values assessed for *F. avenaceum*, *F. poae* and *F. tricinctum* were maximal at ϴ ≥ 25°C, regardless of the a_w_. For *F. langsethiae*, the maximum r values were observed at ϴ ≥ 25°C and low a_w_. The r parameter related to *F. poae* showed a dependence on ϴ only. By contrast, *F. graminearum* showed a complex pattern, with a combined effect of ϴ and a_w_ at a_w_ = 0.97 only, together with a strong effect of extreme a_w_ (0.95 and 0.99).

The Vmax parameter significantly increased with a_w_ levels (p-value < 0.05), with maximal values at high a_w_, for all the ϴ or all species (Figure 3). Temperatures below 20°C were shown to considerably and negatively affect the Vmax parameter for the whole set of studied species.

As appeared in Figure 3, regardless of the considered species, the results showed an increase in τ, and a decrease in K and/or r and Vmax rates under challenging conditions. In any case, growth was promoted by a_w_ ⩾ 97%.

#### INTERSPECIFIC DIVERSITY HAS ONLY A SLIGHT EFFECT ON GROWTH PARAMETERS, COMPARED TO ENVIRONMENTAL FACTORS

The above results demonstrated different growth responses between *Fusarium* species. However, how growth variability is distributed between species, strains and environmental factors (a_w_, ϴ) needs further investigation. The variance partitioning represented in the Figure 4 showed that the studied factors (a_w_, ϴ, species and their interactions) explained 89%, 77% and 73% of the total variance of τ, K and Vmax parameters, respectively. The remaining 11%, 23% and 27% were explained by the residual. A different pattern was observed for r with 52% of the total variance explained by the residual against only 48% by the factors studied. The effect of environmental factors (a_w_, ϴ) and their interaction (a_w_ x ϴ) on the total variance of each growth parameter was predominant and highly significant (p-value < 0.05, see Supplementary Table 2). Nevertheless, the respective contribution of a_w_, ϴ and their interaction (a_w_ x ϴ) to the total variance depends heavily on the measured growth parameter. On the other hand, the species factor alone and its interactions with a_w_ and ϴ explained only 3%, 11%, 8% and 12% of the total variance of τ, K, r and Vmax respectively; the species factor alone was never significant except for K (see Supplementary Table 2).

**Figure 4:**
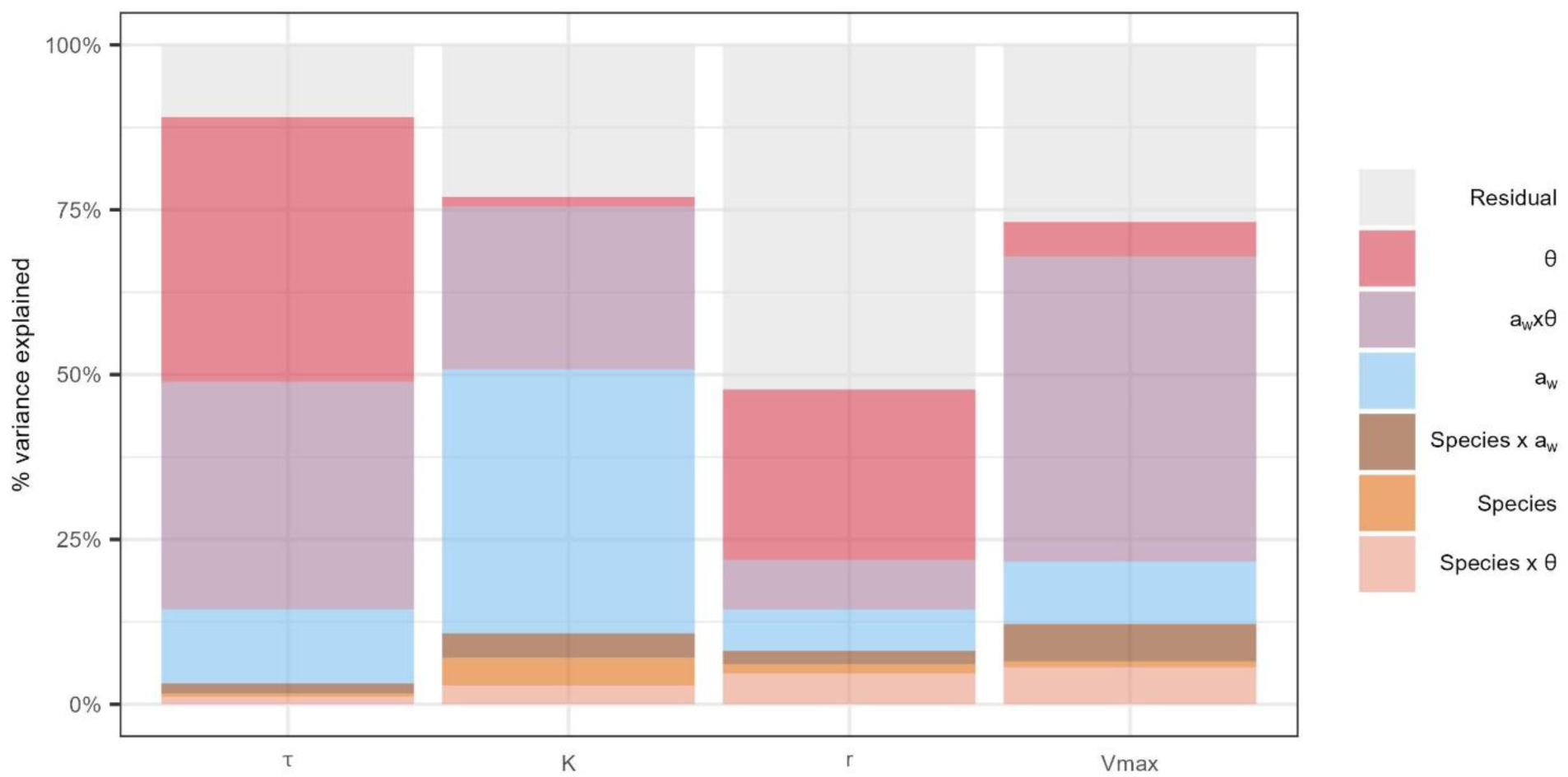
Variance partitioning of growth parameters (τ, K, r and Vmax) according to species, environmental factors (a_w_ and ϴ) and their interactions. The percentage of variance explained (y-axis) by different factors (in colours) for each growth parameter (x-axis) was estimated on grown replicates only. Environmental factors (a_w_ and ϴ), their interaction (a_w_ x ϴ), species factor and its interactions with environmental factors (Species x a_w_ and Species x ϴ) are represented in colours. The strain factor was nested within the species factor.

Our data indicated that ϴ and a_w_ x ϴ interaction factors were the main contributors to the variance of the time to reach inflection point (τ), explaining 40% and 34% of the total variance, respectively, whereas the contribution of a_w_ factor alone was limited to 11% (Figure 4, see Supplementary Table 2).

The variance of the carriage capacity (K) was explained for 40% and 25% by a_w_ and a_w_ x ϴ interaction, respectively. The ϴ factor alone contributed only very slightly (less than 2%) to the K variance (Figure 4, see Supplementary Table 2).

The main factor contributing to the relative growth rate (r) variance was ϴ with 26% of the variance explained, followed by a_w_ and a_w_ x ϴ factors that accounted for 6% and 8%, respectively (Figure 4, see Supplementary Table 2). But, as mentioned above, the residual contributed to half of the variance.

The maximum growth rate (Vmax) variance was shown to considerably depend on a_w_ x ϴ interaction that explained 46% of the variance, while a_w_ and ϴ factors alone accounted only for 9% and 5%, respectively (Figure 4, see Supplementary Table 2).

Altogether, the results reported in Figure 4 allowed to identify some trends related to the impact of environmental factors and their interaction on considered growth parameters. K parameter was mainly affected by a_w_. τ and r were mainly dependent on ϴ, while a_w_ x ϴ interaction contributed importantly to the variance of all parameters.

Surprisingly, species factor and its interactions with abiotic factors contributed only to a low percentage of the variance regardless of the growth parameters. However, as strain factor was nested within the species factor, the interspecific and intraspecific effects on variance were not distinguishable.

#### INTRASPECIFIC DIVERSITY AFFECTS MORE GROWTH PARAMETERS THAN INTERSPECIFIC DIVERSITY

The Figure 5 reports the contribution of both intraspecific (strain factor) and interspecific (species factor) variability to the variance of the growth parameters for each of the a_w_ and ϴ combinations. For a_w_ values higher than 0,97, the intraspecific diversity accounted for more than 45% of the total variance, while interspecific diversity contributed for 30%, regardless of the environmental conditions.

**Figure 5:**
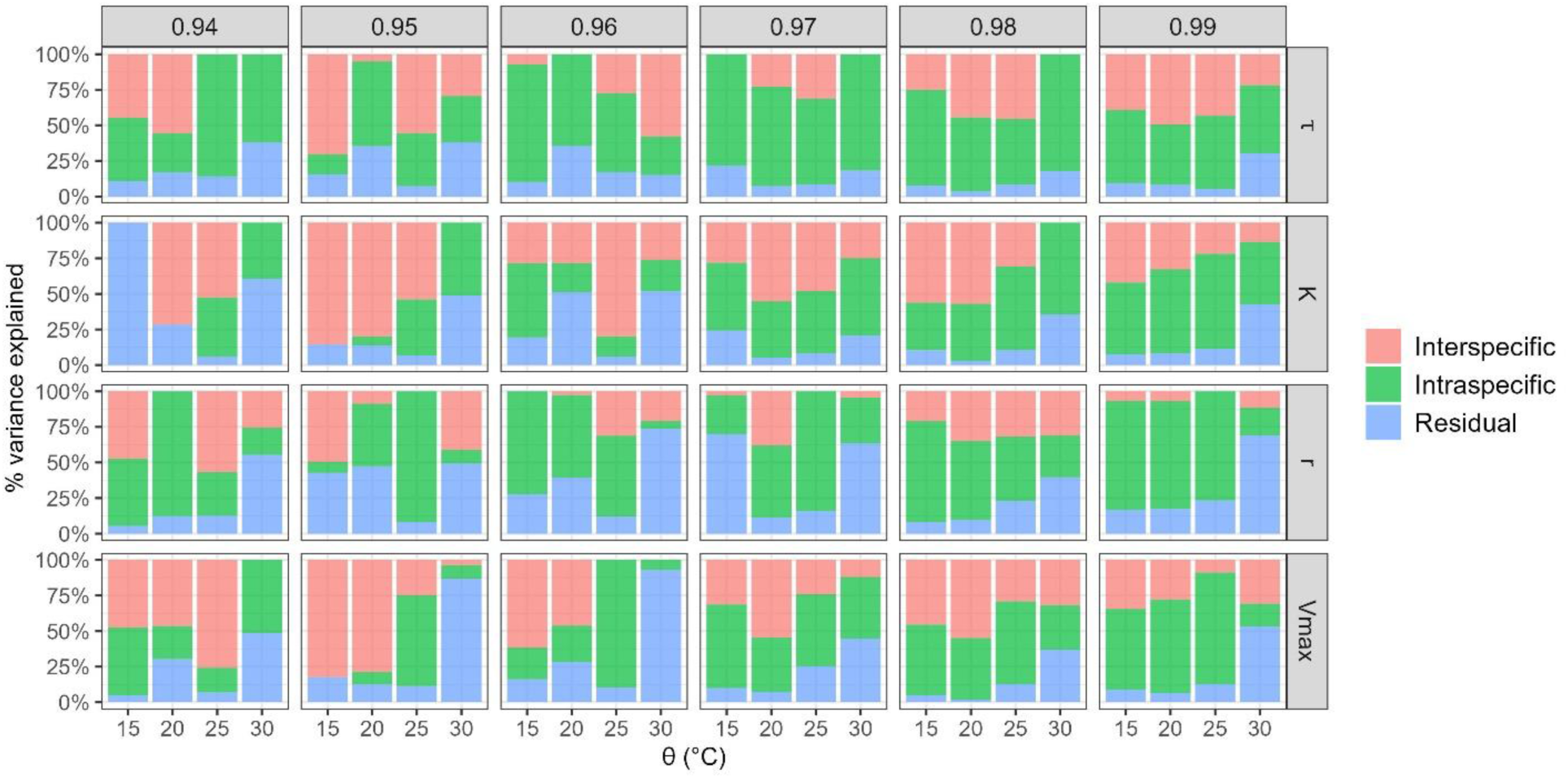
Variance partitioning of growth parameters (τ, K, r and Vmax) according to intraspecific and interspecific diversity. Four different ϴ (15, 20, 25 and 30°C, x-axis) and six levels of a_w_ (0.94, 0.95, 0.96, 0.97, 0.98 and 0.99, in column) were tested to assess the effect of interspecific and intraspecific diversity on the total variance (y-axis).

In contrast, for a_w_ values lower than 0.97, intraspecific variability contributed less to the variance of growth parameters and the effect of the species factor appeared to be more pronounced than the strain factor. For example, 82% and 79% of the Vmax variance was respectively explained by interspecific variability at a_w_ = 0.95 and ϴ = 15°C or ϴ = 20°C (Figure 5).

The residual variance, not explained by the species or the strain, contributed nearly to 24% of the total variance, this percentage increasing in the less favourable growth conditions (15°C and 30°C). In some cases, the residual effect was even stronger than intra- or interspecific effect. For instance, the whole variance of the K parameter was explained by residuals at ϴ = 15°C and a_w_ = 0.94 (Figure 5). However, as mentioned before, estimating the K parameter becomes challenging under severe abiotic constraints, such as a_w_ = 15°C and a_w_ = 0.94 because of a high τ value, preventing the plateau from being reached during the experimentation.

#### INTRASPECIFIC EFFECT IS DIFFERENT BETWEEN SPECIES

Previous results showed a greater effect of the intraspecific diversity on growth parameters, compared to interspecific diversity. The distribution of variability in growth parameters was further studied, taking into account both strain and environmental factors (a_w_, ϴ) and their interaction (a_w_ x ϴ).

Intraspecific variance of growth parameters is reported on Figure 6 for each studied *Fusarium* species. It clearly appeared that this variance was significantly impacted by ϴ, a_w_ and a_w_ x ϴ interaction and that the contribution of the previous factors differed according to the species. Regardless of the species or the studied growth parameter, the proportion of variance explained by the strain factor and its interactions with abiotic factors never exceeded 40%.

**Figure 6:**
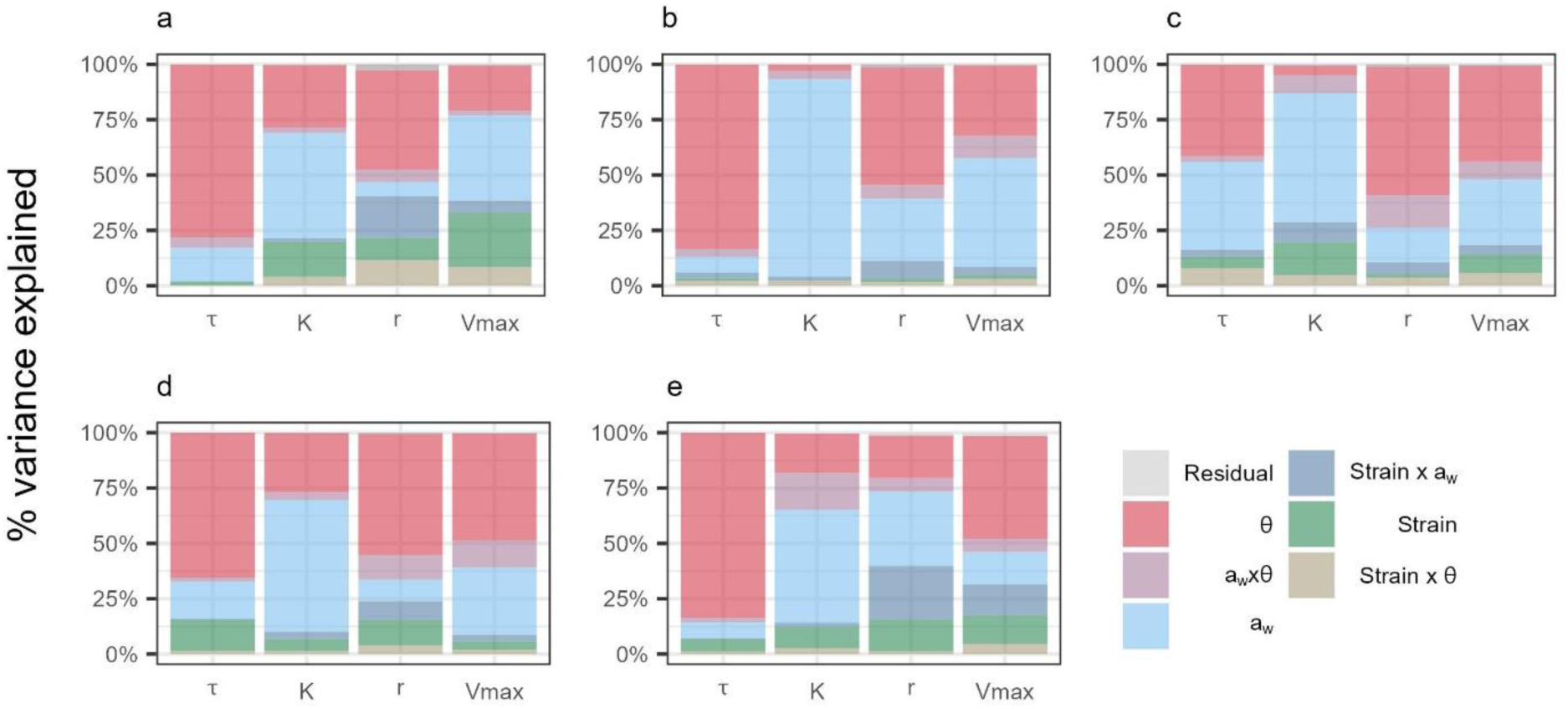
Variance partitioning of growth parameters (τ, K, r and Vmax) for the five *Fusarium* species, according to strain, environmental factors and their interactions. The percentage of variance (y-axis) explained by different factors (in colours) for each *Fusarium* species studied: (a) *F. graminearum*, (b) *F. avenaceum*, (c) *F. langsethiae*, (d) *F. poae* and (e) *F. tricinctum*. Environmental factors (a_w_ and ϴ), their interaction (a_w_ x ϴ), strain factor and its interactions with environmental factors (Strain x a_w_ and Strain x ϴ) are represented in colours.

*F. avenaceum* growth parameters variance strongly depended on environmental factors and their interaction. Indeed, these factors accounted for 94%, 96%, 87% and 91% of the variance of τ, K, r and Vmax variances, respectively (Figure 6b, see Supplementary Table 3). Moreover, the strain factor and interactions contributed only to 6%, 4%, 11% and 8% of the variance (Figure 6b, see Supplementary Table 3).

In the case of *F. graminearum*, environmental factors (a_w_ and ϴ) and their interaction (a_w_ x ϴ) accounted for 98%, 78%, 57% and 61% of the total variance of τ, K, r and Vmax parameters, respectively, while the strain factor and its interactions contributed to 2%, 21%, 40% and 38% (Figure 6a, see Supplementary Table 4). This predominant effect of a_w_, ϴ and a_w_ x ϴ was particularly evident for τ parameter. By contrast, the strain factor and interactions with a_w_ and ϴ contributed to a higher percentage of the variance for r and Vmax parameters.

When growth parameters of *F. langsethiae* followed globally the same pattern, *i.e.* a_w_, ϴ and a_w_ x ϴ strongly contributing to the variance, regardless of the growth parameter (83%, 71%, 88% and 81% for τ, K, r and Vmax, respectively; Figure 6c, Supplementary Table 5), the strain factor and its interactions accounted for a large part of the variance. This was particularly the case for K and Vmax, for which 29% and 18% of their variance was explained by the strain factor and its interactions, respectively (Figure 6c, see Supplementary Table 5).

Considering *F. poae,* environmental factors and their interaction contributed to 84%, 90%, 75% and 91% of the variance of τ, K, r and Vmax parameters, respectively, underlining a small contribution of strain factor and its interactions (Figure 6d, see Supplementary Table 6). Indeed, these factors never explained more than 24% of the total variance, regardless of the parameter.

For *F. tricinctum*, a_w_, ϴ and a_w_ x ϴ contributed to 93%, 85%, 59% and 67% of the variance of τ, K, r and Vmax, respectively, whereas strain factor and its interactions with a_w_ and ϴ accounted for 7%, 14%, 40% and 31% (Figure 6e, see Supplementary Table 7). The predominant effect of environmental factors and their interaction was particularly observed for τ while r and Vmax parameters were also strongly influenced by the strain factor and its interactions.

Overall, environmental factors and their interaction on growth parameters were characterized by a higher impact than the strain factor regardless of the studied species. However the weight of intraspecific variation was different between *Fusarium* species. Indeed, the intraspecific effect was particularly pronounced on growth parameters of *F. graminearum* and *F. tricinctum* while *F. langsethiae* and *F. poae* were less impacted. On the other hand, the intraspecific effect had almost no effect on growth parameters of *F. avenaceum*.

### INFLUENCE OF ENVIRONMENTAL CONDITIONS ON MYCOTOXIN PRODUCTION

#### THE CAPACITY OF SPECIES TO PRODUCE MYCOTOXIN IS DIFFERENTIALLY IMPACTED BY A_W_ AND ϴ FACTORS

The Figure 7 showed that our experimental conditions allowed the production of measurable amounts of mycotoxins by each of the studied species and strains for a_w_ values higher than 0.95 with the exception of some strains of *F. avenaceum*, *F. graminearum*, *F. poae* and *F. tricinctum*, grown at 25°C or 30°C. All strains of *F. graminearum* were able to produce TCTB but with a large intraspecific variability; for example, at ϴ = 25°C and a_w_ = 0.99, *F. graminearum* strains produced between 24 µg.mL^-1^ and 20128 µg.mL^-1^ of TCTB (see Supplementary Figure 3). The amounts of quantified mycotoxins at a_w_ 0.99 indicated that *F. graminearum* produced more TCTB as the ϴ raised (Figure 7). Only the 30°C temperature allowed the production of TCTB at a_w_ = 0.97 and 0.95, the amount strongly decreasing at low a_w_ (Figure 7).

**Figure 7:**
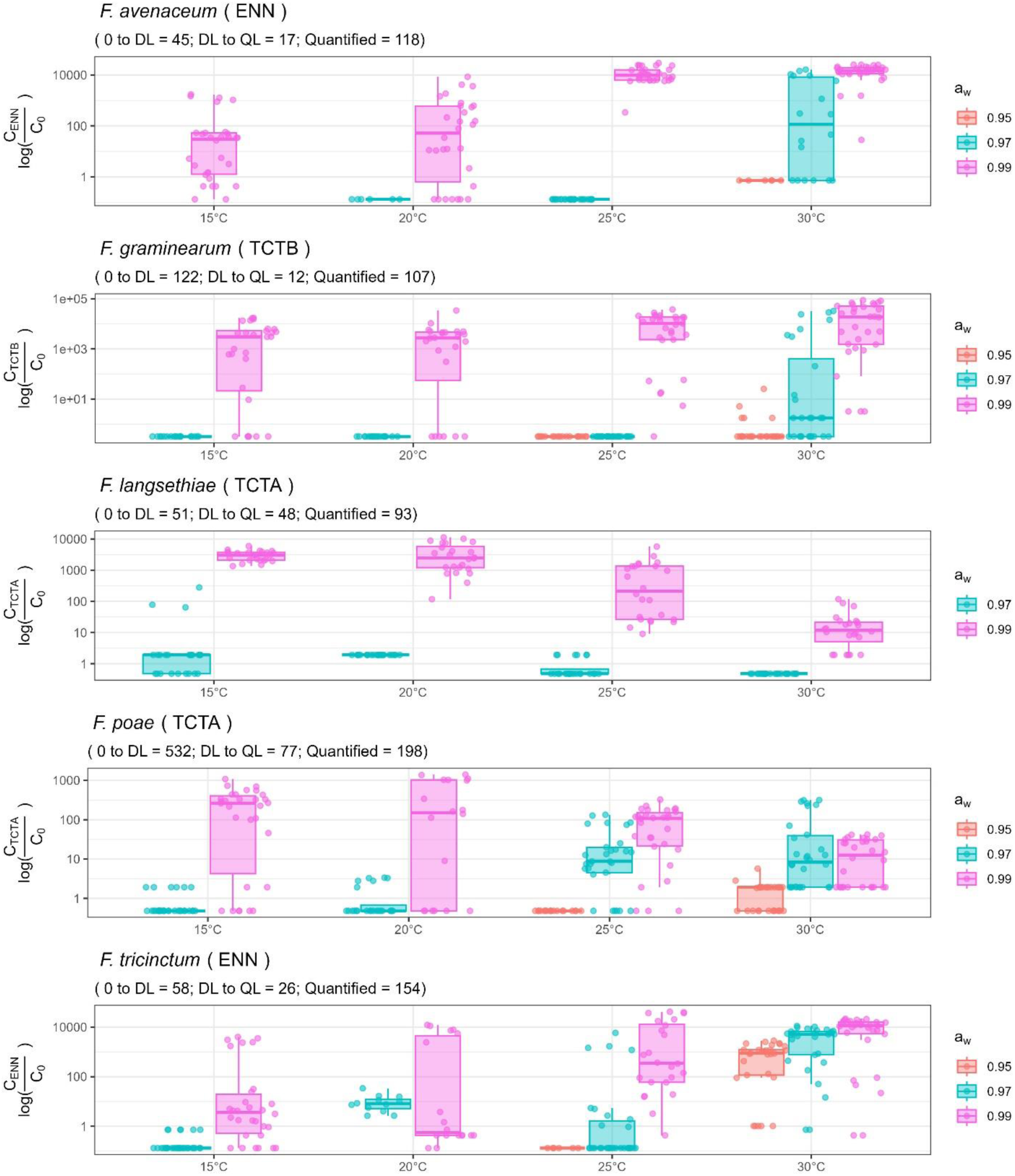
Mycotoxin production by the five *Fusarium* species under environmental variation. Each *Fusarium* species produced a specific class of mycotoxins which are quantified (ENN for *F. avenaceum* in box 1, TCTB for *F. graminearum* in box 2, TCTA for *F. langsethiae* and *F. poae* in boxes 3 and 4, respectively, and ENN for *F. tricinctum* in box 5). Mycotoxins produced (log-transformed, y-axis) were analysed at four ϴ (15, 20, 25 and 30°C, x-axis) and three a_w_ levels (0.95, 0.97 and 0.99, in colours). Each dot represents one replicate and the boxplots show the intraspecific diversity in terms of mycotoxin production within each *Fusarium* species. 0 to DL corresponds to the number of grown replicates in which the mycotoxins were not detected (concentration below the limit of detection (DL). DL to QL corresponds to the number of grown replicates in which the mycotoxins were detected but not quantified because they were below the lower limit of quantification (QL). QL corresponds to the number of grown replicates where the mycotoxins were quantified and shown in these boxplots.

All strains of *F. avenaceum* and *F. tricinctum* produced enniatins in the experimental conditions used throughout this study, the produced amounts varying according to the strain. ENNB (representing 90% and 60% of total enniatins for *F. avenaceum* and *F. tricinctum*, respectively) and ENNB1 (9% and 32%) were the predominant quantified mycotoxins, whatever the ϴ or a_w_ (see Supplementary Figure 3). These two species tended to produce more ENN as the ϴ raised (Figure 7). The highest levels of ENN were quantified at a_w_ = 0.99, 25°C and 30°C. Some ENNs were also quantified in cultures of *F. avenaceum* at a_w_ = 0.97 but only at 30°C (Figure 7). In contrast, the conditions allowing the production of ENNs by *F. tricinctum* were less stringent since measurable amounts of ENNs were observed at a_w_ = 0.97 under 20, 25 and 30°C, and even at a_w_ = 0.95 and 30°C.

Regarding *F. poae*, this species was able to produce mycotoxins belonging to three families (see Supplementary Figure 3) : TCTB (FX), TCTA (DAS and T-2) and ENN (ENNB and ENNB1). A significant correlation (⍴ = 0.8, p-value < 0.05) was observed between TCTB and TCTA, but none with ENN (for example, between TCTA and ENN, ⍴ = 0.33, p-value = 0.38). Indeed, the conditions promoting the production of ENN differed from that of TCTs. DAS was the sole mycotoxin produced above the limit of quantification (QL) by the five *F. poae* strains at all ϴ. When the a_w_ was set at 0.99, the highest production of DAS was observed at low ϴ (15°C and 20°C, Figure 7), while at a_w_ = 0.97 and 0.95, the amount of DAS increased with ϴ (a_w_ = 0.97 at 20, 25 and 30°C ; and a_w_ = 0.95 at 30°C).

All *F. langsethiae* strains were shown to produce TCTA, with the exception of the I508 strain. This strain was also characterized by a weak growth compared to the other strains (data not shown). T-2 was the major produced mycotoxin (78% of the total TCTA), while DAS (14% of TCTA) and HT-2 (8% of TCTA) were quantified in smaller amounts. T-2, HT-2 and DAS were detected at a_w_ = 0.99 under all the tested temperature conditions, the highest amounts being observed at low ϴ (15 and 20°C, Figure 7). At a_w_ 0.97, a low level of TCTA was quantified at 15°C only.

#### PROBABILITY OF MYCOTOXIN PRODUCTION ACCORDING TO ENVIRONMENTAL FACTORS AND SPECIES

Previous results showed that the five *Fusarium* species studied have different abilities for mycotoxin production. The frequency of measurable mycotoxins levels was studied in the subset of samples incubated at a_w_ 0.97 and 0.99 and ϴ = 15°C, 20°C, 25°C and 30°C, thus amounting 699 measurements over 989 grown replicates (see Supplementary Figure 4). Mycotoxins were detected in 71% of the total inoculated wells. Only 57% of *F. graminearum* samples contained mycotoxins, compared to 74% for the four other species. Fitting mycotoxin frequency to the logistic regression model indicated that ϴ, a_w_ and *Fusarium* species factors had significant effects, as well as their interactions (Figure 8).

**Figure 8:**
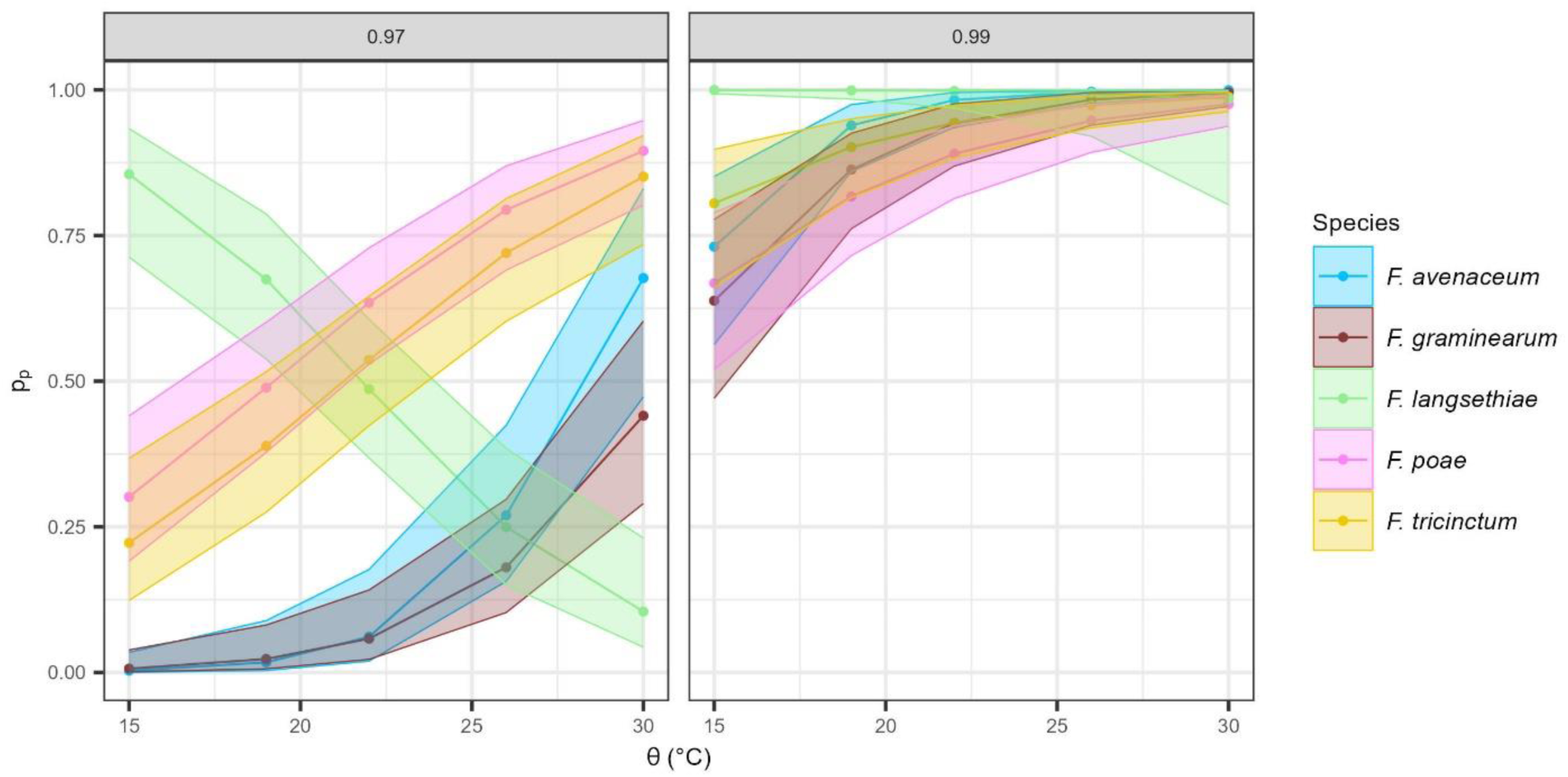
Probability of mycotoxin production for the five *Fusarium* species, according to environmental variation. The probability of mycotoxin production (y-axis) was estimated by the logistic regression model (logit(p_p_) = β_0_ + (β_1_ + β_1i_)·ϴ_i_ + β_2i_·a_wi_ + β_12i·_a_wi_·ϴ_i_ + β_3j_·fs_j_ + β_ij_a_wi_ fs_j_) against four ϴ (15, 20, 25 and 30°C, on the x-axis) and at two levels of a_w_ (0.97 and 0.99, in column). Each dot represents the probability of mycotoxin production of the five *Fusarium* strains for each species. Confidence intervals (95%) are represented by shading around the curves. Data for a_w_ = 0.95 are not shown because very few grown replicates were able to produce mycotoxins above the quantification threshold.

The probability of mycotoxin production (p_p_) for each studied species is reported in Figure 8. As a general pattern, p_p_ increased with a_w_ and ϴ. At an a_w_ of 0.99, the probability that a species produces measurable levels of mycotoxins is high, especially for ϴ > 20°C. However, as previously reported for growth, *F. langsethiae* displayed a distinct behaviour compared to the other species. At a_w_ of 0.97, the probability of mycotoxin production by this species strongly decreased with the ϴ (Figure 8). When comparing the five species at a_w_ = 0.97, it appeared that *F. poae* and *F. tricinctum* were characterized by the highest p_p_.

A censored 2-way ANOVA was performed using the whole set of mycotoxin data acquired in the present study (see Supplementary Figure 5). This allowed to clearly determine differences in the effects of ϴ and a_w_ on the production of mycotoxins by each of the studied *Fusarium* species. A_w_ was shown to significantly modulate the mycotoxin production, regardless of the species, while ϴ had a significant impact only for *F. graminearum, F. langsethiae* and *F. tricinctum.* This last species was the only one affected for its mycotoxin production by the a_w_ x ϴ interaction (see Supplementary Figure 5, see Supplementary Table 8).

#### CORRELATION BETWEEN ENVIRONMENTAL CONDITIONS, MYCOTOXINS AND GROWTH PARAMETERS

Correlations between growth parameters and mycotoxin production were analysed, considering the studied environmental factors (a_w_, ϴ) and their interaction (a_w_ x ϴ). All data (ranks of growth parameters and mycotoxin) related to each fungal species were combined in PCA analyses (Figure 9). The first axis represented 50% to 56% of the variance depending on the considered species. For *F. avenaceum, F. graminearum, F. langsethiae* and *F. tricinctum*, the first axis was shown to discriminate samples characterized by an early (τ) and high (Vmax) growth leading to high OD at 14 days, together with high mycotoxin production levels (ρ = 0.39 to 0.57), from late and low growth samples, together with low mycotoxin production levels. The second axis represented 23 to 27% of the variance, leading to a negative correlation between ranks of τ and r (ρ = -0.51). The third axis, contributed to 6 to 15% to the variance. This axis primarily reflected variations in mycotoxin ranks, which tended to be higher in samples grown at high a_w_. The mycotoxin ranks moderately correlated with other growth parameters (ρ = 0.31 for K, -0.36 for τ, and 0.13 for r) and more strongly with OD at 14 days (ρ = 0.48). The previous observations suggest that samples characterized by a delayed growth were associated with a lower production of mycotoxins. Overall, results presented in Figure 9 clearly show that mycotoxin production is correlated to growth parameters, regardless of the species, and differentially driven by environmental factors (a_w_, ϴ). Finally, the growth and the mycotoxin production of *F. graminearum, F. avenaceum* and *F. tricinctum* were correlated and were specifically favoured by ϴ ≥ 25°C.

**Figure 9:**
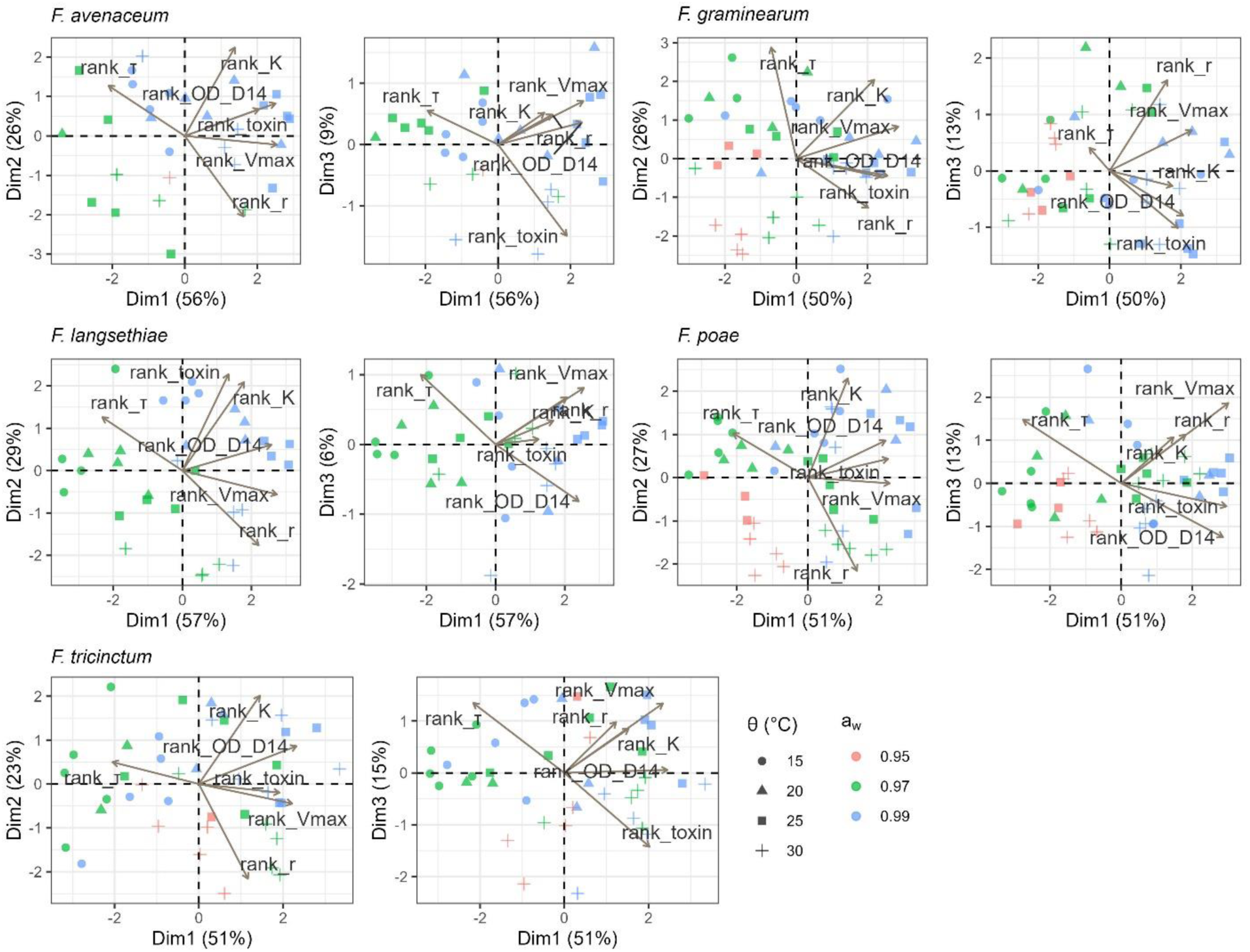
Principal component analysis of mycotoxin and growth parameters (τ, K, r and Vmax) for the five *Fusarium* species. The ranks of growth parameter and mycotoxins for *F. avenaceum*, *F. graminearum*, *F. langsethiae*, *F. poae* and *F. tricinctum* are presented at four ϴ (15, 20, 25 and 30°C) and three a_w_ levels (0.95, 0.97 and 0.99). For each species, first biplot is for (Dim1, Dim2) and second biplot is for (Dim1, Dim3). Each dot corresponds to a strain growing at specific a_w_ and ϴ.

## DISCUSSION

*Fusarium* Head Blight (FHB) is a multi-pathogen disease caused by a consortium of several *Fusarium* species occupying the same niche. A more comprehensive understanding of the intricacies of such multiple infections is essential to predict the outcome in terms of disease development but also mycotoxin production. Climate is certainly the first driving force that shape *Fusarium* spp. occurrence in the field (Perrone *et al*. 2020). Using *in vitro* studies, we investigated the behaviour of major *Fusarium* species under various challenging environmental conditions.

Several studies have already addressed the impact of abiotic constraints on the growth and mycotoxin production of *Fusarium* species (Medina *et al*. 2010; Peter Mshelia *et al*. 2020; Yu *et al*. 2021). But to the best of our knowledge, no comprehensive analysis as the one discussed in the present study has been published before. Indeed, this study is the first one considering five major *Fusarium* species causing FHB and a large range of combined temperature and water activity conditions. In addition, whereas most of the earlier studies have mainly focused on one or a couple of strains, the present one was based on a set of five strains per species. Lastly, the analytical process of our dataset was optimized by the use of innovative statistical approaches.

### OPTIMISING ANALYSES TO FULLY EXPLOIT RELEVANT BIOLOGICAL DATA

Inferring fungal growth rates from optical density (OD) measurements has been already proven to be relevant, either for yeast-like microorganism or for filamentous fungi (Boixel *et al*. 2019; Hameed *et al*. 2024). In this paper, the OD readings were directly used as raw data, generating a large dataset that allowed robust statistical analyses for modelling growth and estimating time-course growth parameters for each *Fusarium* species. Thus, the observed growth curves were consistent with theoretical growth patterns, displaying the four expected phases; lag, exponential, deceleration and stationary (see Supplementary Figure 1), despite hyphal growth and the possible production of secondary metabolites that could affect OD measurements. The measurements were repeatable across replicates and revealed different responses to environmental conditions. Such high-throughput phenotyping experiments, with a large number of strains and replicates, rapidly become time consuming, generate a lot of waste, can be quite onerous and require an optimal valorisation of acquired data (Heidorn 2008; Zamir 2013). This optimal valorisation was a secondary objective of the present work. To achieve this objective, appropriate statistical methods were used to ensure optimum preservation of the relevant information. For example, generalized additive models (GAM) have enabled accurate mapping of the OD of the culture medium, improving the sensitivity and specificity of fungal growth and estimated growth curves, even at very low values of OD (Wood 2017). Furthermore, instead of discarding wells lacking growth/mycotoxin production, we used logistic regression models which provided valuable insights into the intrinsic dynamics of these wells. As far as we know, this was the first time where growth and mycotoxin production were expressed as probability. Such insights are particularly relevant in the context of *Fusarium* Head Blight (FHB), where the presence or absence of species in the field carries significant implications beyond mere quantitative considerations. Moreover, in the analysis of mycotoxins, we decided to not discard values falling below detection/quantitation limits (observed in 46% of the samples) and to not substitute them with arbitrary values, a practice discouraged by Shoari and Dubé (2018) for example. By employing methods tailored to handle censored data, we were able to explore a broader spectrum of environmental conditions.

### DIFFERENTIAL RESPONSES OF *FUSARIUM* SPECIES UNDER FLUCTUATING ENVIRONMENTAL CONDITIONS

Growth and mycotoxin production responses to changing environmental conditions were shown to differ according to the studied *Fusarium* species. Our results indicated that *F. graminearum* was the most affected in its growth ability by challenging environmental conditions, suggesting that this species could display a lower phenotyping robustness than the other species. Our data agree with previous reports indicating highest growth rates at a_w_ = 0 .99 and ϴ = 25°C for *F. graminearum* (Hope *et al*. 2005).

Consistently, the growth probability assessed for *F. graminearum* was lower than that of the other *Fusarium* species, except at 25°C. This observation could be the result of different ecophysiological requirements of *Fusarium* species particularly for growth initiation. Indeed, in the literature, 25°C is often considered to be the optimum temperature for *F. graminearum* germination (Manstretta *et al*. 2016).

We showed that *F. langsethiae* exhibited a different behaviour compared to *F. graminearum*, *F. avenaceum*, *F. poae* and *F. tricinctum*. Notably, the highest probabilities of growth and mycotoxin production for this species were observed at 15°C.

We have also observed that combined environmental factors, water activity (a_w_) and temperature (ϴ), impacted significantly both the growth dynamics and mycotoxin production, regardless of the species. Our findings showed contrasted responses of τ and K parameters, suggesting that the growth rate is more affected by ϴ, while a_w_ has a greater influence on final growth. Verheecke-Vaessen *et al*. (2022) observed similar results using three *F. langsethiae* strains, demonstrating that the lag phase was predominantly impacted by ϴ. Additionally, the growth of *F. graminearum* and *F. culmorum* was reported as being strongly affected by ϴ variations (Hope *et al*. 2005). Our data also demonstrated that mycotoxin production is strongly affected by a_w_ levels and ϴ, which is consistent with previous studies (Kokkonen *et al*. 2010; Medina and Magan 2011). The latter authors observed a major effect of a_w_ on T-2 and HT-2 mycotoxin production by *F. langsethiae*. We specifically observed that the production of mycotoxins by *F. graminearum* was more restrained in challenging conditions than that of the other species, as described by Hope *et al*. (2005).

Moreover, we highlighted the correlation between growth and end-point mycotoxin production, emphasising the importance of the mycotoxin production dynamics, as described by Verheecke-Vaessen *et al*. (2022). Further analysis investigating the production of mycotoxin by *Fusarium* species for longer incubation times could allow clarifying whether low amounts of mycotoxin result from a delay in production or an inhibition induced by challenging environmental conditions (Hope *et al*. 2005).

### EXPLORING INTRASPECIFIC DIVERSITY OF *FUSARIUM* SPECIES

An essential and thus far under-investigated aim of this study was to consider the intraspecific diversity within five major *Fusarium* species causing FHB. In less challenging growth conditions, we found that intraspecific diversity accounted for a high proportion of the variance compared to interspecific diversity. However, when environmental factors (a_w_, ϴ) were included in the analysis, it became clear that their combined effects were governing growth variation at the species and strain levels. This indicates that while intraspecific diversity has a significant impact on *Fusarium* species behaviours, its effect is negligible compared to abiotic factors, particularly in more challenging environments.

Besides, our findings illustrate the extent of the phenotypic responses to abiotic constraints exhibited by each species, suggesting the existence of genotypes that are likely to be better adapted. Our data corroborate previous reports indicating that *Fusarium* species and different strains within a same species exhibit varying degrees of resilience to environmental fluctuations and that such a diversity should absolutely be considered in initiatives aiming to predict the impact of abiotic factors on FHB. They support the conclusion of Ma *et al*. (2013) that has highlighted the difficulty of predicting FHB risk due to the occurrence of populations of distinct strains in the field.

### BEYOND ECOPHYSIOLOGY OF *FUSARIUM* SPECIES: ANTICIPATE THEIR FUTURE DISTRIBUTION UNDER CLIMATE CHANGE

Climate change is expected to strongly impact the distribution of *Fusarium* species in Europe (Parikka *et al*. 2012) and could exacerbate the damages related to FHB including the contamination of grains with mycotoxins (Moretti *et al*. 2019). We confirmed that environmental conditions are likely to be the main driver of the predominance of one *Fusarium* species over the others. Interestingly, our results based on *in vitro* investigations support previous results from field surveys that have shown the occurrence of shifts in *Fusarium* species distribution. For example, Covarelli *et al*. (2015) confirmed the predominant effect of thermo-hygrometric conditions in the replacement of *F. graminearum* by *F. avenaceum* and *F. poae* between 2009 and 2010. Other studies have also shown that the occurrence of *F. graminearum* is promoted by moderately warm summers whereas dry and hot climates are more favourable to *F. poae* or *F. avenaceum* (Van Der Lee *et al*. 2015; Banik *et al*. 2018; Nogueira *et al*. 2018; Valverde-Bogantes *et al*. 2020). Predicting the distribution of FHB species and the associated mycotoxin risk is therefore essential to limit disease outbreaks and ensure feed and food safety.

Battilani *et al*. (2013) have established a mechanistic weather-driven model based on the life cycle of *Aspergillus flavus* to predict field contamination by aflatoxins. They emphasised the importance of including ecophysiological parameters of fungal species in the models. A similar conclusion was raised by Van Der Fels-Klerx *et al*. (2012) who have attempted to model climate change effects on wheat phenology and DON contamination in wheat cultivated in North West Europe by 2040. The previous authors have used various climatic and agronomic variables to calculate a principle empirical model underlying the biology of *Fusarium* infection and DON production. The present study highlighted intraspecific and interspecific ecophysiological properties of *Fusarium* species under environmental variations. These experimental data could be used as variables to calibrate the model parameters and improve the prediction of FHB risk and mycotoxin contamination.

In order to draw a comprehensive pattern of response to abiotic conditions, we investigated the ecophysiology of *Fusarium* species, one by one. Nevertheless, in the field, a cohort of *Fusarium* species co-occurs during FHB infection (Arseniuk *et al*. 1999). The composition of this *Fusarium* community changes over time and location, and depends both on environmental and agronomic factors (Karlsson *et al*. 2021), influencing the abundance of one species compared to another (Boutigny *et al*. 2019).

Furthermore, in addition to the ecophysiological characteristics of each species, their capacity to compete is also a key criterion governing their occurrence. As an example, *F. graminearum* is described as being one of the most competitive species when interacting with other *Fusarium* species on wheat spikes (Audenaert *et al*. 2013; Siou *et al*. 2015). However, our results suggest that such a dominance could be questioned under challenging conditions. It is now imperative to investigate the ecophysiology of different *Fusarium* species in interaction in order to better understand the influence of abiotic factors on the competition outcome and to construct a more accurate representation of what happens in the field.

## CONCLUSIONS

The present study highlighted differences in ecophysiological properties of the main *Fusarium* species causing FHB in response to the combined effects of temperature and water activity. Considering five species and five strains per species grown under 24 different environments, acquired data have allowed delivering new and significant insights: (1) growth and mycotoxin production of *F. graminearum* and *F. avenaceum* was favoured by high temperatures and high water activity, (2) lower water activity levels were more conducive for *F. poae* and *F. tricinctum*, (3) *F. langsethiae* exhibited a specific fitness for the colder temperature (4) beyond contrasted behaviours between species, a large phenotypic diversity within species was also highlighted. While our data offer a comprehensive foundation for enhancing predictive models of FHB, further studies encompassing more intricate aspects, including fungal interactions and environmental factors influencing the entire life cycle, are essential to achieve greater precision.

## MATERIALS AND METHODS

### GROWTH AND MYCOTOXIN PRODUCTION STRAIN COLLECTION

Five strains of five *Fusarium* species *F. avenaceum, F. graminearum, F. langsethiae*, *F. poae* and *F. tricinctum* were used within this study (see Supplementary Table 1). These 25 strains (isolated from wheat, maize or barley grains) are registered in public biological resource centers or are available at the MycSA laboratory.

### CULTURE MEDIA

All strains were inoculated in standardized sucrose synthetic liquid medium (MS) containing 20 g.L^-1^ of sucrose, 0.5 g.L^-1^ KH_2_PO_4_, 0.6 g.L^-1^ K_2_HPO_4_, 0.017 g.L^-1^ MgSO_4_, 1 g.L^-1^ (NH_4_)_2_SO_4_, and 0.1 mL.L^-1^ Vogel mineral salts solution. Glycerol (Sigma-Aldrich, Saint-Louis, USA) was added to liquid medium according to the Norrish equation to adjust the a_w_ values between 0.94 and 0.99. The resulting a_w_ was controlled using an A_w_-meter (TESTO 650, Lenzkirch, Germany). A total of 24 conditions of incubation were considered: temperatures (ϴ) 15, 20, 25 and 30°C combined to a_w_ = 0.94, 0.95, 0.96, 0.97, 0.98 and 0.99.

### SPORE PRODUCTION

Fungal strains were grown on Potato Dextrose Agar (PDA, Difco, Le-Pont-de-Claix, France) solid medium for seven days. For each strain, the spore solution was prepared by inoculating seven agar plugs in 75 mL of carboxymethylcellulose (CMC, Sigma-Aldrich, Saint-Louis, USA) liquid medium. The cultures were incubated three days in the dark with shaking (180 rpm) at 25°C before being filtered (40 µm filter) and centrifuged at 4800g for 20 minutes. The supernatants were discarded and the spore pellets were resuspended in 4 mL of MS. The spores were counted on a Malassez cell and diluted to reach a final concentration of 1,33.10^6^ spores.mL^-1^. Batches of spores were kept in 50% glycerol (50:50, v/v : glycerol:MS medium) at -80°C until inoculation.

### INOCULATION AND STUDY OF FUNGAL GROWTH

Batches of spores were thawed the day of inoculation. Each strain was inoculated in six replicates at a final concentration of 1,66.10^4^ spores.mL^-1^ in 200 µL of MS liquid medium in 96-well plates. Half of the wells of each plate were filled with blanks (non-inoculated wells, with MS at the correct a_w_) to avoid contamination between the wells and to allow data to be corrected from background between experiments if necessary (see “Estimation of the absorption of the plates” section). Plates were sealed with Breath-easy sealing membrane (Diversified Biotech, Boston) to avoid evaporation and incubated in the dark at the selected ϴ. Optical density (OD) was measured in each well every 24 hours, five time points, at five different spatial positions in each well, using a spectrophotometer Infinite M200 pro (Tecan, Grödig, Austria) set at λ = 630 nm until the growth stationary phase was observed.

### STUDY OF MYCOTOXIN PRODUCTION

The same inoculation procedure as described in the previous section was used with the following modifications: 24-well plates were used with a final volume of 2 mL of MS and six wells per plate were used as controls (non-inoculated wells, with MS at the correct a_w_). Twelve conditions of incubation were performed : 15, 20, 25 and 30°C combined to a_w_ = 0.95, 0.97 and 0.99. At the end-point time, *i.e.* after 14 days of incubation, the cultures were centrifuged (20 minutes at 4800g, ambient temperature). Supernatants were kept at -20°C until mycotoxin extraction and the mycelia were rinsed with water, conserved at -80°C before freeze-drying. Mycotoxins were extracted using 3 mL of ethyl acetate as extraction solvent. Briefly, the samples were shaken for 5 minutes and centrifuged 10 minutes at 4800g (ambient temperature). Upper phase was recovered and evaporated with a SpeedVac Concentrator SPD210 (Asheville, USA) at 55°C. Then, 200 µL of methanol/water (50:50, v/v) were added, and after homogenization with a Vortex mixer, extracts were filtered through a filter syringe (Acrodisc 13 mm mini spike with 0.2 µm wwPTFE; VWR International, Puerto Rico) and kept at -20°C until analysis.

### MYCOTOXINS ANALYSIS

A Vanquish UHPLC (ThermoScientific, Bremen, Germany) system was used to separate mycotoxins (Type B and A trichothecenes and Enniatins) on a reversed-phase column Thermo Accucore aQ C18 (2.1 x 100 mm, 2.6 µm; ThermoScientific, Lithuania) maintained at 40°C. Mobile phase consisted of water/methanol (98:2, v/v) (eluent A) and methanol/water (98:2, v/v) (eluent B) both containing 5 mM of ammonium acetate and 0.1% (v:v) acetic acid. Gradient elution was as follows: 2% of B for 0.5 min, 2 to 98% B in 4.5 min, 98% B for 4.7 min, 98 to 2% for 0.1 min and 2% B for 2.2 min. The flow was kept at 0.4 mL.min^-1^ and the injection volume was 2 μL. Mycotoxins were detected using a Q Exactive Focus mass spectrometer (ThermoScientific, Bremen, Germany). Heated electrospray ionization source was operated in positive mode. Full scan spectra were acquired at a 70k resolving power in the 150-1000 *m/z* range. The Orbitrap analyzer was *m/z*-calibrated each week. Nitrogen was used as the sheath and auxiliary gas. The main MS parameters were set as follows: sheath gas flow rate, 50 psi; auxiliary gas flow rate, 13 a.u.; sweep gas flow rate, 0 a.u.; spray voltage, 3.5 kV; capillary temperature, 320°C; S lens RF level 50%; auxiliary gas heater temperature, 350°C. External calibration was performed at 5, 10, 25, 50, 100, 250, 400 and 500 µg.L^-1^ to quantify TCTB (Deoxynivalenol (DON), 3- and 15-ADON as 15-acetyldeoxynivalenol (15-ADON), Zearalenone (ZEA), Nivalenol (NIV) and Fusarenon-X (FX)), TCTA (Diacetoxyscirpenol (DAS), T-2 and HT-2), enniatins (ENNA, A1, B and B1) and beauvericin (BEA). Mix standard of TCTB, TCTA and ZEA in acetonitrile (10 µg.mL^-1^) was purchased from Romer labs (Tulin, Austria) and mix of ENN (A, A1, B, B1) and BEA in methanol (10 µg.mL^-1^) from LIBIOS (Vindry-sur-Turdine, France).

Some mycotoxins were not detected (concentration below limit of detection (DL)), others were detected but not quantified (concentration below the lower limit of quantitation (QL)).

The limit of quantification (QL) was estimated at 5 µg.L^-1^ for ENN quantification and at 25 µg.L^-1^ for TCTB and TCTA.

Mycotoxin production was calculated by the sum of the different compounds within each group: TCTB (DON + 15-ADON), ENN (ENNA + ENNA1 + ENNB + ENNB1) and TCTA (DAS + T-2 + HT-2) and expressed as final concentration in µg.mL^-1^.

## DATA ANALYSIS

All statistical analyses were performed with R Studio software version 4.3.1 (2023-06-16).

### ESTIMATION OF THE ABSORPTION OF THE PLATES

To avoid potential biases due to the light absorption by plates and to liquid media interference, 48 non-inoculated wells by plate (MS at the correct a_w_) were used to assess and model in both spatially and temporally terms, the blank absorption. At each time point (every 24 hours), a spatial Generalized Additive Model (GAM) was applied using the *mgcv* package (Wood 2017) to interpolate the optical density (OD) values across the entire plate. GAM uses splines, which are smooth functions composed of piecewise polynomials, to relate the X and Y positions of each well to the OD values. GAM was chosen for its flexibility, smoothness, and anisotropic nature (ability to vary in different directions, see Supplementary Figure 1). After estimation, the GAM-predicted OD values attributed to the microplates (OD_well_) were subtracted from the measured OD values in each inoculated well, yielding:

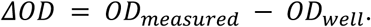

### FITTING OF THE GROWTH CURVES

For each well, ΔOD was fitted for each day (D) using a sigmoidal growth curve, following the equation reported below:

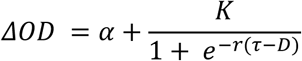

α is a parameter describing the well-specific noise of the measurements due to medium absorption and has no biological meaning, while K, r and τ are parameters characterising the kinetics of ΔOD influenced by the combination of ϴ and a_w_. K represents the carriage coefficient of the environment and it is expressed without unit, r is the relative growth rate in OD_unit_.days^-1^ and τ is the time of the inflection point of the curve expressed in days (see Supplementary Figure 2). The growth rate at inflexion time (Vmax expressed in OD_unit_.days^-1^) is obtained as rK/4. We then employed the *drm* function from the *drc* package to fine-tune the curves (Ritz *et al*. 2015).

### MODELLING THE PROBABILITY OF GROWTH

Depending on the experimental conditions, some strains did not grow at all in some replicates, while the other replicates showed measurable growth. Growth probability was first approximated for each species by calculating the growth frequency (f_g_, Figure 1) of the six replicates for each strain. The probability of growth (p_g_, Figure 2) was then modelled, using a logistic regression.

The logistic regression method assumes a linear relation between the logit of the growth probability (p_g_) and the covariates; temperature (ϴ), water activity (a_w_), and *Fusarium* species (fs):

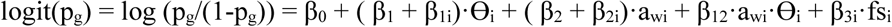

where β_1_ and β_2_ correspond to the effect of ϴ and a_w_, respectively; β_3i_ corresponds to the effect of *Fusarium* species i; β_1i_ (and β_2i_) corresponds to the interaction between *Fusarium* species and ϴ (respectively a_w_); and β_12_ represents the interaction between ϴ and a_w_.

The optimal model was selected among all possible combinations of covariates using the Bayesian Information Criterion (BIC). The logistic regression method is valuable as it offers an analytical approach to model the growth probability. The *glm* function base was used to adjust the logistic regression model.

### VARIANCE PARTITIONING ANALYSIS

To investigate the influence of each factor on each parameter (K, r, τ and Vmax), a type-III ANOVA, incorporating ϴ, a_w_, *Fusarium* species, and their respective interactions as covariates was applied. The strain was nested within the *Fusarium* species factor. The results of the ANOVA analysis were used to decompose the variance explained by each of the factors and their interactions. This analysis was done using the *anova* function, together with the *lmer* function of the *nlme* package.

The proportions of variance explained by the species and the strain for each parameter, at each level of ϴ and a_w_, were compared, using the *partvar* function of the *cati* package (Taudiere and Violle 2016). Effect of the intraspecific diversity was investigated by running an ANOVA within each species on each parameter, with ϴ and a_w_ as covariates.

### MODELLING THE PROBABILITY OF MYCOTOXIN PRODUCTION

Mycotoxin production was not dynamically measured since the procedure used in the present study was destructive (refer to the section “Study of mycotoxin production”).

Depending on experimental conditions, some species/strains/replicates were shown to produce quantifiable amounts of mycotoxins while others did not. Similarly as what was done for growth probability, logistic regressions were applied to model the probability of mycotoxin production.

The logistic regression method assumes a linear correlation between the logit of the mycotoxin production probability (p_p_) and the covariates; temperature (ϴ), water activity (a_w_), and *Fusarium* species (fs). The optimal model was selected among all possible combinations of covariates using the BIC.

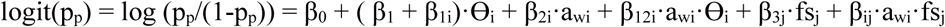

where β_1_ corresponds to the effect of ϴ, β_1i_ (and β_2i_) corresponds to the interaction between species and ϴ (respectively a_w_), β_12i_ represents the interaction between ϴ and a_w_, β_3j_ corresponds to the effect of *Fusarium* species j, and β_ij_ corresponds to the interactions between a_w_ and the *Fusarium* species j.

However, for *F. langsethiae*, numerical issues were faced since the a_w_ factor induced a separation problem (when a covariate completely explains an outcome; Albert and Anderson 1984) where all samples exhibited growth at a_w_ = 0.99. This separation led to huge confidence intervals and infinite parameters in logistic regression. To overcome this issue, the method proposed by Heinze and Schemper (2002) was applied. Briefly, this method aims to maximize a penalised likelihood to get a finite estimate of the parameters. It is available in the *logistf* package.

### EFFECT OF ENVIRONMENTAL CONDITIONS ON THE MYCOTOXIN PRODUCTION

In some grown replicates, mycotoxin levels were below the QL (left-censored observations). Common practices for handling left-censored observations involve either discarding values below the quantitation limit or substituting them with an arbitrary value (*i.e.* half of the quantitation threshold). However, these approaches may i) discard valuable information from the original data and/or ii) introduce biases in summary statistical calculations (Shoari *et al*. 2016; Shoari and Dubé 2018).

Therefore, for each species and associated mycotoxin group, a censored version of a two-way ANOVA was done. This version of the ANOVA allows to test differences between groups on a quantitative variable, while retaining left-censored observations. The effect of ϴ, a_w_ and their interaction was tested and the model having the best BIC was selected. The function from the *NADA2* package (v1.1.; Julian and Helsel 2024) was used.

### CORRELATIONS BETWEEN MYCOTOXINS PRODUCTION AND GROWTH PARAMETERS

The correlation between mycotoxin production and growth parameters was investigated using a Principal Component Analysis (PCA) on the ranks of both growth parameters and mycotoxins quantification, for each species and strain. To mitigate scale effects resulting from the different mycotoxins, the ranks of mycotoxins (*i.e.* ordered mycotoxins based on their values relative to the dataset) were computed within each group of mycotoxins (TCTB, TCTA and ENNs). To take into account the fact that under challenging growth conditions, some samples did not finish their growth at 14 days, the OD at 14 days was included in the analysis when mycotoxins were measured. PCA on the ranks allowed to keep many values without introducing biases caused by arbitrary data substitution.

## ACKNOWLEDGEMENTS

We thank Sylvain Chéreau for the initial development of the LC-MS method. We also thank Maider Abadie for her English supervision.

## DATA AVAILABILITY

Data and scripts are available in the data INRAE repository:

